# Cycloserine for Treatment of Multidrug-Resistant Tuberculosis in China: A Retrospective Observational Study

**DOI:** 10.1101/307124

**Authors:** Yang Li, Fei Wang, Limin Wu, Min Zhu, Guiqing He, Feng Sun, Qihui Liu, Xiaomeng Wang, Wenhong Zhang

## Abstract

**Objectives:** Cycloserine is crucial in multidrug-resistant tuberculosis (MDR-TB) treatment. Although extensive research has been carried out on MDR-TB, most researchers have not treated cycloserine in much detail. Therefore, we evaluate the efficacy and safety of cycloserine and seek to clarify the role of cycloserine for treatment of simple MDR-TB, pre-extensively drug-resistant tuberculosis (pre-XDR-TB), and extensively drug-resistant tuberculosis (XDR-TB).

**Patients and methods:** A retrospective observational study was performed in China. We determined the treatment outcome as the primary outcome for 144 cycloserine-treated and 181 cycloserine-nontreated patients according to the definitions of WHO. The proportion of patients with sputum-culture conversion and the frequency of adverse drug reactions related to cycloserine were assessed as well.

**Results:** Among 325 MDR-TB patients, 144 were treated with cycloserine and 100 (69.4%) out of 144 successfully completed treatment. Compared with patients in non-cycloserine group, the hazard ratio of any unfavorable treatment outcome was 0.53 (95%CI: 0.35-0.81, *P*=0.003). Culture conversion rate at the intensive phase was similar whether cycloserine was administered or not (*P*=0.703). Of the 144 patients treated with cycloserine, a total of 16 (11.1%) patients experienced side-effects related to cycloserine, including 7 patients who discontinued cycloserine permanently.

**Conclusions:** Cycloserine could be an attractive agent to treat MDR-TB. Its safety profile warrants use in the most of MDR-TB cases. Cycloserine significantly improved the chance of favorable outcome for patients with simple MDR-TB but not pre-XDR-TB and XDR-TB. More aggressive regimens might be required for pre-XDR-TB or XDR-TB patients.

## Introduction

Tuberculosis (TB) has been a continuing threat throughout the ages. Since the early 1990s, the global outbreaks of multidrug-resistant tuberculosis (MDR-TB), defined as tuberculosis (TB) caused by organisms that are resistant to rifampicin and isoniazid, have been reported and it is generally accepted that resistance to these two potent anti-TB agents is associated with an increased probability of catastrophic treatment costs and poorer treatment outcomes. Furthermore, extensively drug-resistant tuberculosis (XDR-TB), defined as MDR-TB plus resistance to a fluoroquinolone and an injectable second-line drug, has recently emerged and threaten the public health on a worldwide scale(1). In 2016, there were an estimated 490000 new cases of MDR-TB and globally the second highest number of drug-resistant TB (DR-TB) cases came from China. The latest data from the World Health Organization (WHO) reported a treatment success rates of 54% for MDR-TB and only 30% for XDR-TB(2).

Cycloserine, a cyclic analogue to D-analogue, could target alanine racemase and D-alanine ligase, thus blocking the formation of bacterial cell wall(3). Cycloserine has been introduced in tuberculosis therapy since the late 1950s(4). Years later, Sommer *et al* found its potential for therapeutic intervention for chronic pulmonary tuberculosis(5). However, the neurological toxicity of cycloserine has been concerning clinicians and limiting its use widely. An earlier report described the neurological adverse effects of cycloserine that it would induce symptomatic seizures in approximately 10% patients(6). With more effective drugs like rifampicin discovered, cycloserine has been applicable only in the treatment of apparent or proved drug-resistant tuberculosis.

To implement tuberculosis control, WHO has published treatment guidelines for drug-resistant TB in 1997 for the first time and cycloserine has been suggested since then as it shares no cross-resistance with other agents and might be valuable to prevent resistance to other active drugs(7). Cycloserine was classified as a Group 4 oral bacteriostatic second-line medication in the 2008 recommendations(8). A study from Turkey in 2011 reported their experience in treating MDR-TB that the overall success rate of treatment achieved to 77% with the use of intensive regimens which included cycloserine(9). Meanwhile, WHO guidelines for the treatment of drug-resistant TB was updated and suggested a stronger association with cure of cycloserine than para-aminosalicylic acid (PAS)(10). In 2016, WHO regrouped the anti-TB agents and cycloserine was included in Group C (other core second-line agents) together with ethionamide (or prothionamide), linezolid and clofazimine. Generally, two or more of Group C agents are to be included when designing the core MDR-TB treatment regimen and ethionamide (or prothionamide) and cycloserine are supposed to be selected in preference to linezolid and clofazimine(2), which means that cycloserine would be in MDR-TB treatment’s starting line-up. However, the clinical studies which mainly focused on cycloserine, particular in East Asia patients, are scarce, as the use of cycloserine was not approved in China until recently. In addition, the role of cycloserine in the treatment of XDR-TB or pre-XDR-TB (defined as resistance to isoniazid and rifampicin plus any fluoroquinolone or one of the injectable drugs) is not clear-out. To light of the uncertainties, we aimed to provide sufficient details to evaluate the efficacy, tolerability and safety of cycloserine in MDR/pre-XDR/XDR-TB treatment using a sizable cohort of patients with MDR-TB from China.

## Patients and methods

### Study design and procedures

This retrospectively cohort study was performed at two hospitals located in Zhejiang Province, China initiated by the Zhejiang Disease Control and Prevention Center (CDC) who has set up routine drug resistance monitoring for TB since 1999(11). Patients aged above 18 who were diagnosed with active MDR-TB were recruited consecutively during March 2012 through December 2015 to obtain full follow-up information. Positive culture for *Mycobacterial tuberculosis* and resistance to isoniazid and rifampicin proven by drug-susceptibility testing were required at enrolment. Furthermore, the participants were included when their treatment therapies were adapted to WHO recommendations (2016 version). Patients were excluded when met any of the following criteria: (1) positive for HIV test; (2) history of seizure disorder, mental depression, or severe anxiety; (3) decline to participate in this study.

The following information were collected: sociodemographic characteristics, indicators of severity (symptoms and radiologic findings), previous treatment, drug-resistant profiles, and background treatment regimen. Culture and sputum conversion and chest X-rays were performed periodically for treatment outcomes evaluation. Moreover, adverse drug reactions were monitored and promptly managed during the entire treatment course.

Approval for collection of data was provided by the ethics committees of Zhejiang CDC. All patients provided written informed consent.

### Definitions

MDR-TB was defined as tuberculosis caused by a strain as *M. tuberculosis* that was resistant to at least isoniazid and rifampicin. XDR-TB was MDR-TB that was also resistant to the fluoroquinolones and any of second-line injectable drugs (capreomycin, kanamycin and amikacin). Pre-XDR-TB was MDR-TB that was resistant to either a fluoroquinolone or a second-line injectable drug, but not both. In the present study, we use the term simple MDR-TB to refer to those with resistance to just isoniazid and rifampicin and complicated MDR-TB to refer to those with additional resistance beyond isoniazid and rifampicin including pre-XDR-TB and XDR-TB. Standard treatment outcome definitions were applied according to the definitions and reporting framework for TB from WHO in 2013(12). Cured was defined as treatment completed without evidence of failure and three or more consecutive cultures were negative after the intensive phase. If bacteriological results were lacking (*i.e.* fewer than three cultures performed), the case was defined as treatment completed. Treatment failure was defined as treatment terminated or need for permanent regimen change of at least two anti-TB drugs because of lack of conversion by the end of the intensive phase, or bacteriological reversion in the continuation phase after conversion to negative, or adverse drug reaction. Default was defined as interruption of treatment for at least 2 months not meeting the criteria for failure. This study used the following brief outcomes: favorable outcome was defined as cured and treatment completion; unfavorable outcome was defined as any failure, default or death while on treatment.

When assessing the adverse drug reaction, we distinguished two types of side-effects: major side-effects and minor side-effects(13). The formed referred to any adverse reactions that resulted in temporary or permanent discontinuation of anti-TB drugs, while the later referred to those that only required dose adjustment and/or addition of concomitant treatment.

### Drug susceptibility testing

Sputum culture on Löwenstein-Jensen medium or MGIT 960 were applied routinely. Phenotypic drug susceptibility testing to two first-line drugs (rifampicin, isoniazid) and two second-line drugs (ofloxacin and kanamycin) was performed from the first positive *Mycobacterial tuberculosis* culture with the use of the proportion method and the result was compared with the standardised strains. The critical drug concentrations of rifampicin, isoniazid, ofloxacin, and kanamycin were 40, 0.2, 2, and 30 μg/ml respectively (14).

### Data management and statistical analysis

The clinical data were collected through questionnaires and medical records by trained health workers. For the analysis, patients were divided into two cohorts according to the presence or absence of cycloserine in the background regimen (Cycloserine cohort *versus* Non-cycloserine cohort). Continuous variables were calculated as mean with standard deviation (SD) and median with interquartile range and were further compared by Mann-Whitney *U* test. Categorical data were presented as numbers (percentage) and were compared with the use of χ^2 test.

The primary outcome was the proportion of favorable treatment in each treatment cohort. Secondary outcome included the efficacy of cycloserine measured by the proportion of conversion within the intensive phase and safety and tolerability of cycloserine measured by the frequency of major and minor reactions.

For the primary outcome, all patients’ treatment outcomes were identified according to the definitions described above by two clinicians blinded for the background regimen. And the proportion of each treatment outcome for two cohort were calculated. Considering the potential confounders, we further investigated the effect of cycloserine upon the treatment outcome by using a Cox proportional-hazards model. Furthermore, we did the specific subgroup analyses of patients with different drug resistance patterns.

Two-tailed *P* value of less than 0.05 was considered statistically significant. All statistical calculations and analyses in this study were performed with the use of SPSS Statistics, version 22.0 (IBM).

## Results

### Study population

Enrolment of patients began in March 2012 and the follow-up for the last patient was performed by December 2017. A total of 582 patients were assessed for eligibility. 241 patients were excluded because their background regimens were not adapted to WHO recommendations (2016 versions) and 11 patients were excluded because the strains from their isolates were identified as nontuberculous mycobacteria. Moreover, three patients with HIV positive and two patients with mental illness in the control group were excluded as well. Consequently, 325 patients confirmed to have an organism resistant to both rifampicin and isoniazid were enrolled, 144 of whom were treated with cycloserine in their background regimen according to WHO guidelines for designated dosages of 500mg or 750mg per day (500mg for 38 patients weight less than 50kg; 750mg for 96 patients more than 50kg). All patients’ background regimen included one of fluoroquinolones and only two patients in cycloserine group had not been treated with aminoglycosides as the initial treatment. Most of demographic and baseline clinical characteristics were comparable among two treatment cohorts except tuberculosis cavity being more frequent in the cycloserine group. The mean age was 42.9 years and approximately 70% of patients were male. 27.4% (89/325) patients were treated with at least one of fluoroquinolones or aminoglycosides more than 30 days before. More details could be found in Table 1.

**Table 1.**
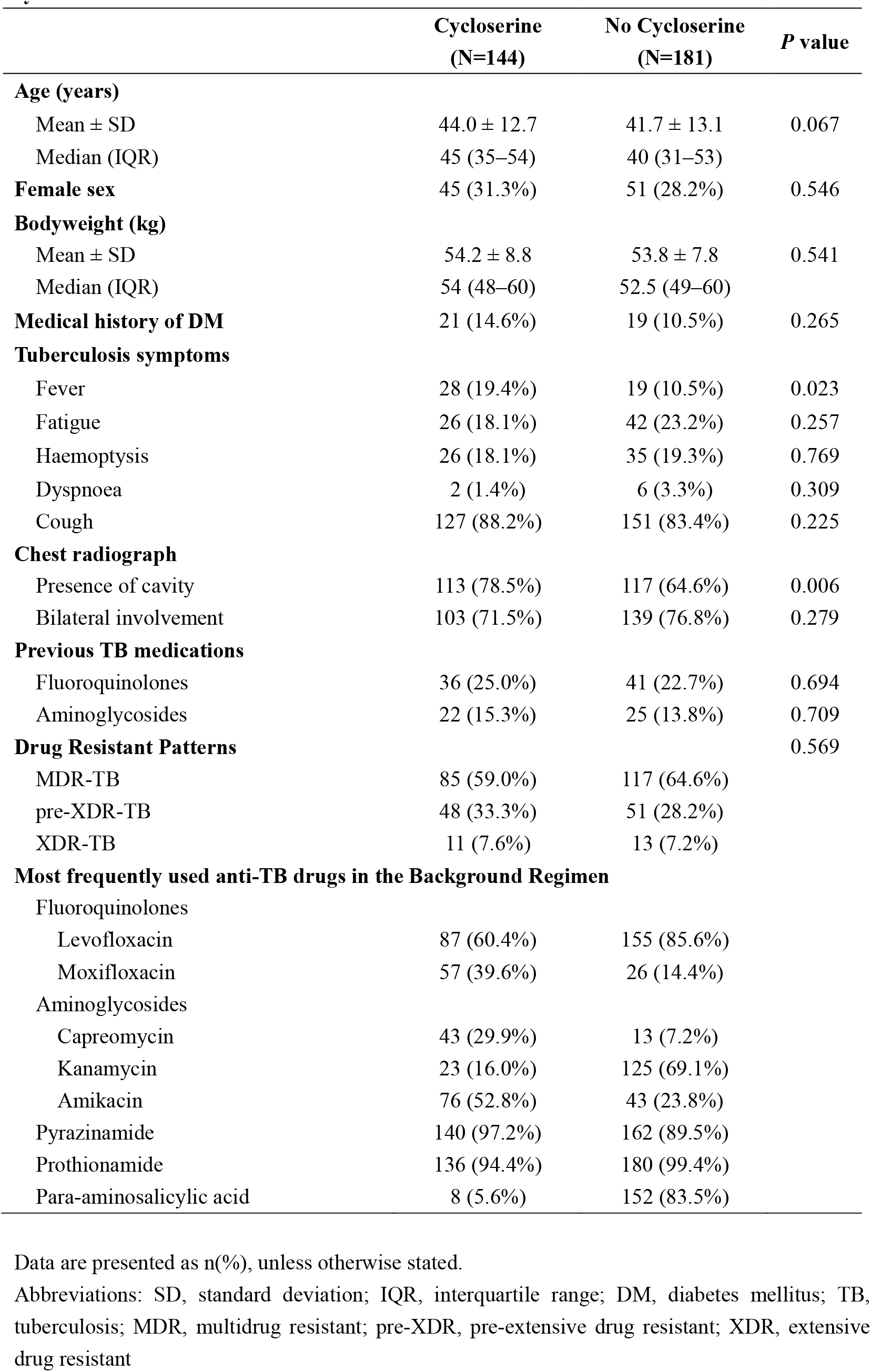
Characteristics of multidrug resistant tuberculosis cases treated with or without cycloserine.

### Treatment outcome

There was a trend approaching a level of significance in treatment outcome: 100 out of 144 (69.4%) cycloserine-treated patients achieved treatment success *versus* 108 out of 181 (59.7%) non-cycloserine-treated patients (χ^2 test, *P*=0.089, Table 2). The absence to sputum conversion at 6 months and severe adverse drug effects resulting in two or more drugs stoppage were the main reason for treatment failure; the relative responsibilities were 35.1% and 43.3% in the cycloserine group and 41.7% and 43.3% in the non-cycloserine group respectively. One patient was complicated by pulmonary infection and died on the eighteenth months of treatment. To reduce confounding bias, a Cox regression analysis was used and suggested that introduction of cycloserine to the standard drug regimen resulted in significantly less risk of unfavorable treatment outcomes (Hazard Ratio, [HR]: 0.58, 95% confidence interval [CI]: 0.35-0.81, *P*=0.003, Table 3).

**Table 2.** Comparison of treatment outcomes of multidrug-resistant/extensively drug resistant tuberculosis cases treated with or without cycloserine.

|  | **Cycloserine**<br>**(N=144)** | **No Cycloserine**<br>**(N=181)** | ***P* value** |
| --- | --- | --- | --- |
| **Treatment Success** | **100 (69.4%)** | **108 (59.7%)** | **0.089** |
| Cure | 94 (65.3%) | 106 (58.6%) |  |
| Treatment Completion | 6 (4.1%) | 2 (1.1%) |  |
| **Treatment Failure** | **37 (25.7%)** | **60 (33.2%)** |  |
| Fail to conversion at 6 months | 13 (9.0%) | 26 (14.4%) |  |
| Reversion | 8 (5.6%) | 9 (5.0%) |  |
| Adverse drug reactions | 16 (11.1%) | 25 (13.8%) |  |
| **Death** | **0 (0.0%)** | **1 (0.5%)** |  |
| **Default** | **7 (4.9%)** | **12 (6.6%)** |  |

**Table 3.** Cox regression analysis of potential independent variables associated with unfavorable treatment outcome in multidrug resistant tuberculosis cases.

| Variables | Univariate Cox Regression |  | Multivariate Cox Regression |  |
| --- | --- | --- | --- | --- |
|  | Crude HR<br>(95%CI) | *P* value | Adjusted HR<br>(95%CI) | *P* value |
| Age ≥ 60 years | 2.23 (1.32–3.66) | 0.003 | 2.61 (1.46–4.65) | 0.001 |
| Cough | 3.11 (1.45–6.68) | 0.004 | 2.34 (1.07–5.11) | 0.033 |
| Fever | 1.83 (1.17–2.87) | 0.008 | 1.57 (0.95–2.61) | 0.079 |
| Bilateral involvement | 1.92 (1.18–3.10) | 0.008 | 1.45 (0.88–2.38) | 0.141 |
| Previous fluoroquinolones treatment* | 1.80 (1.20–2.70) | 0.004 | 1.71 (1.08–2.83) | 0.024 |
| Previous aminoglycosides treatment* | 1.71 (1.10–2.69) | 0.019 | 1.01 (0.58–1.76) | 0.969 |
| Resistance to fluoroquinolones | 2.55 (1.57–4.14) | <0.001 | 1.92 (1.16–3.16) | 0.011 |
| Resistance to aminoglycosides | 2.61 (1.37–4.99) | 0.004 | 1.35 (0.68–2.70) | 0.391 |
| Cycloserine treatment<sup>†</sup> | 0.66 (0.45–0.96) | 0.030 | 0.53 (0.35–0.81) | 0.003 |
| Pyrazinamide treatment<sup>†</sup> | 0.38 (0.22–0.66) | 0.001 | 0.70 (0.20–2.41) | 0.570 |
| Clarithromycin treatment<sup>†</sup> | 2.32 (1.44–3.72) | <0.001 | 1.97 (0.91–4.25) | 0.085 |
| High-dose isoniazid treatment<sup>†</sup> | 2.23 (1.20–4.15) | 0.012 | 0.58 (0.15–2.25) | 0.430 |
| Amoxicillin–clavulanate treatment<sup>†</sup> | 4.11 (1.30–12.99) | 0.016 | 5.07 (1.42–18.17) | 0.013 |

Abbreviations: HR, Hazard Ratio; CI, confidence interval
\* Treated with fluoroquinolones or aminoglycosides more than 30 days before.
<sup>†</sup> Treated with cycloserine, pyrazinamide, clarithromycin, high-dose isoniazid or amoxicillin-clavulanate as the baseline regimen.

### Efficacy end-points assessment

Efficacy was mainly measured by sputum culture conversion and proved to be roughly similar between two groups. There was no difference in the proportion achieving sputum culture conversion at the end of intensive phase (117/144,81.3% *versus* 144/181,79.6%, *P*=0.703) or at the end of treatment (127/144,88.2% *versus* 149/181,82.3%, *P*=0.142) between the cycloserine group and non-cycloserine group. For those who had sputum culture conversion, the mean±SD time to culture conversion in patients treated with cycloserine were longer than those without (90±121 days *versus* 59±61 days, *P*=0.003). With the use of Cox regression analysis, cycloserine also did not accelerate sputum culture conversion (HR: 1.057, 95%CI: 0.814-1.372, *P*=0.679).

### Safety assessment

Overall, 132 of 144 patients (91%) in the cycloserine group and 161 of 181 patients (89%) in the non-cycloserine group had clinically significant adverse drug reactions. The most frequent adverse events were gastrointestinal effects (nausea and vomiting), arthralgia, liver injury and hypokalaemia in both two treatment groups (Figure 1). Among these 132 patients reporting adverse events in the cycloserine group, 37 (28%) experienced major adverse-effects, whereas 95 (72%) patients experienced minor side-effects. Adverse events attributed to cycloserine are shown in Table 4. Side-effects that were possibly or probably related to cycloserine appeared after a median of 71 days (range 10-331 days) of cycloserine treatment. A total of sixteen patients reported seventeen episodes related to cycloserine, including nine patients discontinued cycloserine temporarily or permanently. We observed eight episodes of headache and cycloserine was permanently withdrawn from the treatment regimen in two patients. Moreover, two cases of seizure, one case of depression, and two cases of anxiety were observed, with these events resulting in cycloserine discontinuation within the first six months of treatment. No suicidal ideation was observed.

**Figure 1.**
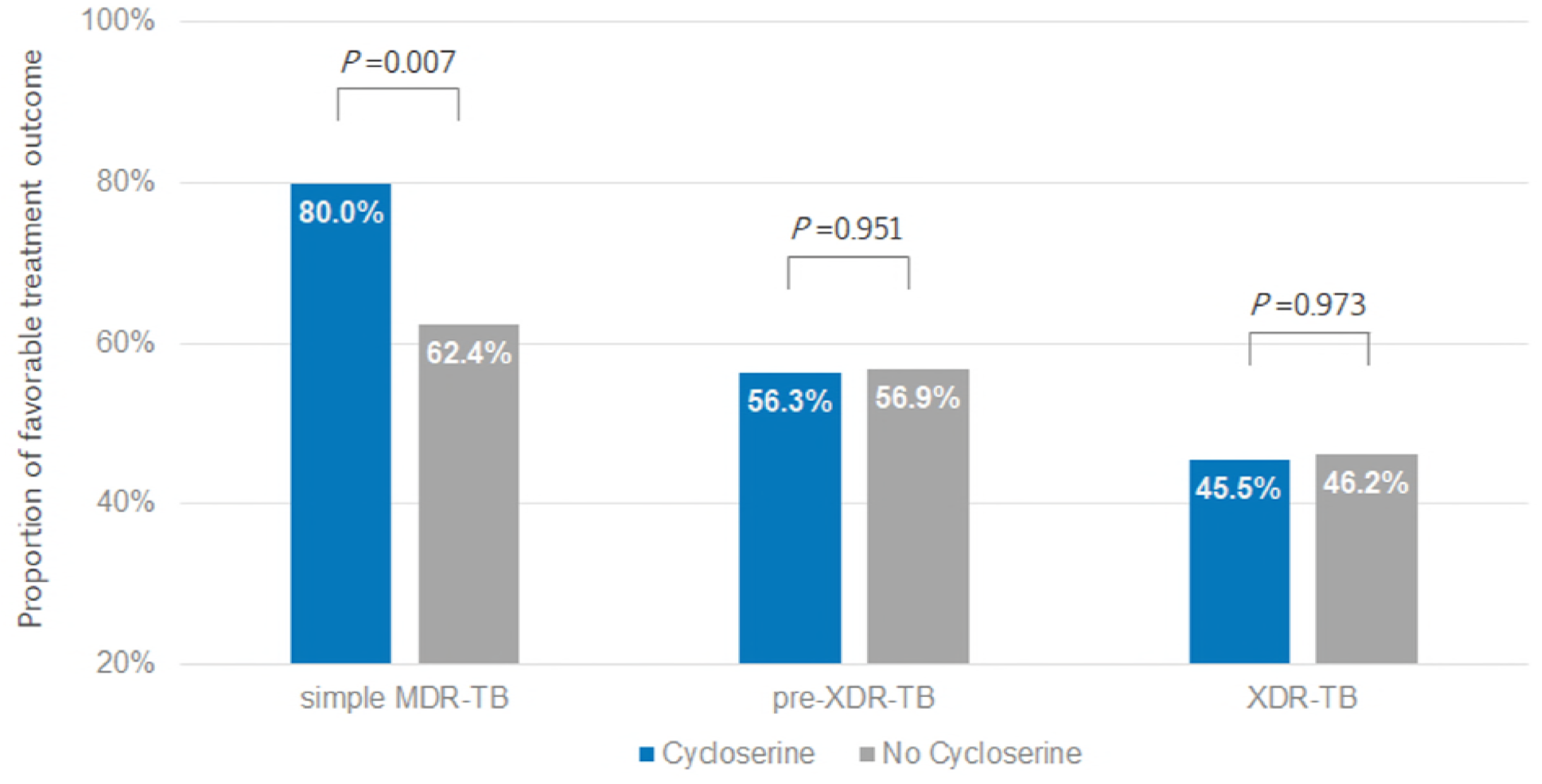
Proportions of favorable treatment outcome, according to the resistance pattern among patients treated with and without cycloserine. Abbreviation: MDR-TB, multidrug resistant tuberculosis; XDR-TB, extensively drug-resistant tuberculosis.

**Table 4.** Side effects associated with cycloserine or requiring to withdraw cycloserine.

| Patient ID | Age (years) | Daily CS doses* | Interval time† (days) | Side effects | Relationship to CS | Doses Adjustment or Stoppage |
| --- | --- | --- | --- | --- | --- | --- |
| P9 | 48 | 500mg | 119 | abdominal distension | Unlikely related | De-escalation to 250mg |
| P80 | 52 | 750mg | 115 | headache | Probably related | temporarily stopped (2 days) |
| P89 | 51 | 500mg | 132 | depression | Probably related | permanently stopped |
| P90 | 39 | 750mg | 10 | seizures | Possibly related | permanently stopped |
| P96 | 28 | 750mg | 164 | headache | Probably related | permanently stopped |
| P106 | 56 | 500mg | 127 | seizures | Possibly related | permanently stopped |
| P109 | 57 | 500mg | 16 | headache | Probably related | No adjustment |
|  |  |  | 16 | peripheral neuropathy | Possibly related | No adjustment |
| P110 | 61 | 500mg | 80 | tremor | Possibly related | permanently stopped |
| P112 | 63 | 500mg | 41 | rash | Possibly related | No adjustment |
| P115 | 45 | 750mg | 331 | headache | Possibly related | De-escalation to 500mg |
| P117 | 49 | 500mg | 50 | headache | Probably related | permanently stopped |
| P119 | 45 | 750mg | 28 | anxiety | Possible related | temporarily stopped (1 month) |
| P123 | 38 | 750mg | 58 | headache | Probably related | No adjustment |
| P124 | 35 | 750mg | 154 | headache | Probably related | De-escalation to 500mg |
| P127 | 37 | 750mg | 71 | headache | Probably related | No adjustment |
| P143 | 39 | 750mg | 21 | anxiety | Probably related | permanently stopped |

\*Daily CS doses refer to the doses in the background regimen.
†Interval time from start of therapy to appearance of side effects (days).
Abbreviation: CS, cycloserine; mg, milligram.

### Treatment outcomes stratified by resistance patterns

The treatment outcomes were further compared between two groups stratified by resistance patterns (Figure 2). Among simple MDR-TB patients, the proportion of treatment success in the cycloserine group was higher than the non-cycloserine group, reaching statistical significance (68/85, 80.0% *versus* 73/117, 62.4%, *P*=0.007). For other strata, the treatment success rate in the cycloserine group was almost similar to patients who were not treated with cycloserine. Or rather, among pre-XDR-TB patients, the proportion achieving favorable outcome was 56.3% (27/48) in those receiving cycloserine *versus* 56.9% (29/51) in those treated without cycloserine (*P*=0.951); and among XDR-TB patients, the proportion of favorable outcome was 45.5% (5/11) and 46.2% (6/13), respectively (*P*=0.973). We also calculated the sputum conversion rate at six months and observed no significant difference between two groups regardless of the resistance patterns (data not shown). Moreover, a downward trend in favorable treatment outcome rate was observed with the increase in the extent of drug resistance in both groups.

**Figure 2.**
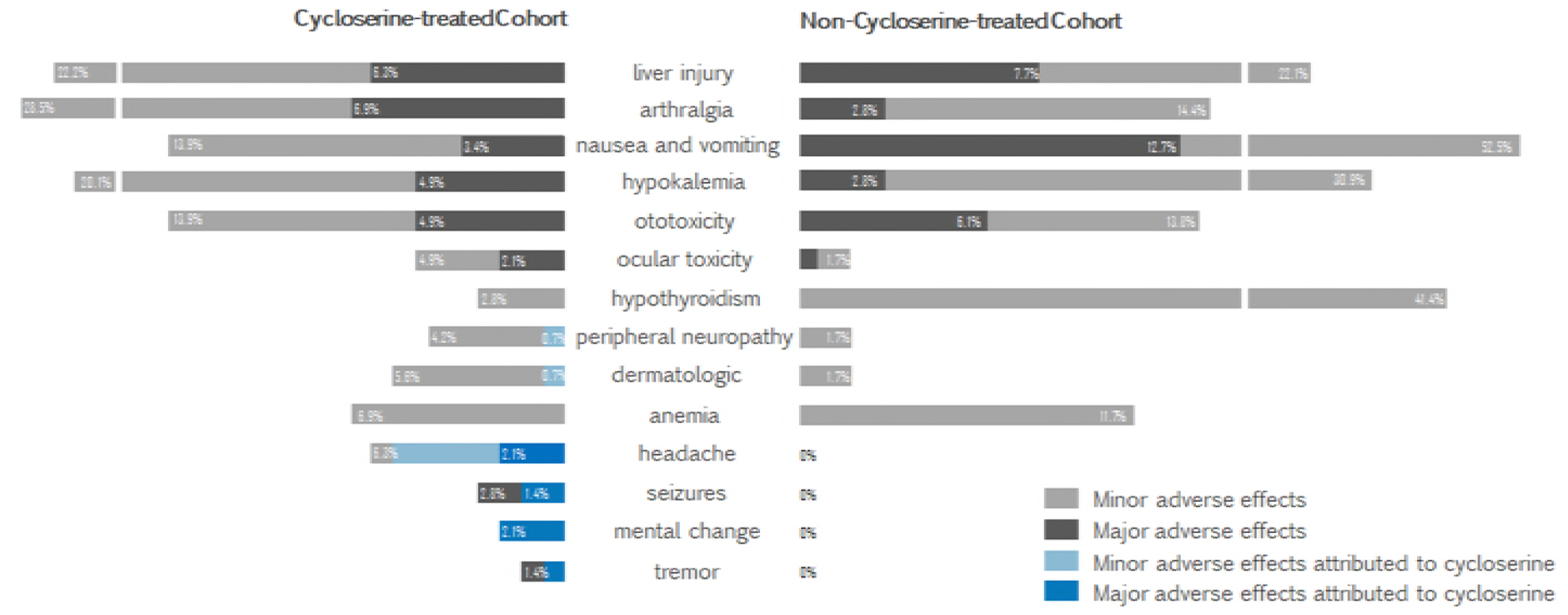
Adverse drug reactions in patients with and without cycloserine treated for multidrug-resistaπt tuberculosis in China. Adverse drug reactions which were associated with cycloserine are marked in sky blue (Minor adverse effects) or navy blue (Major adverse effects).

### Risk factors to unfavorable treatment outcomes

The associations between unfavorable treatment outcomes and each baseline variables were firstly assessed with univariate Cox regression model. Those variables with *P* value < 0.1 would be included into multivariate Cox regression model (Table 3). Using a Cox regression analysis, we found that a significant risk of unfavorable treatment outcomes related to age older than 60 years (HR: 2.61, 95%CI: 1.46-4.65; *P*=0.001), presence of cough before starting treatment (HR: 2.34, 95%CI: 1.07-5.11; *P*=0.033), and previous exposure to fluoroquinolones more than 30 days (HR: 1.71, 95%CI: 1.08-2.83; *P*=0.024) or resistance to fluoroquinolones proven by drug-susceptibility testing (HR: 1.92, 95%CI: 1.16-3.16; *P*=0.011).

## Discussion

The research on tuberculosis treatment has witnessed a clear shift from drug-sensitive tuberculosis to drug-resistant tuberculosis, as more than half of patients with drug-resistant tuberculosis experience treatment failure owing to the weaknesses and intolerability of current treatment regimen for drug-resistant tuberculosis. To improve the treatment outcome, intensified research and innovation, the third pillar of WHO’s Post-2015 Global Tuberculosis Strategy, has been emphasized(15). Safer, easier and shorter treatment regimens is a critical target that clinicians are moving forward to. The arrival of novel drugs like delamanid and bedaquiline has offered fresh opportunities(16)(17) but up to now, there are not enough new drugs to establish an entirely new regimen, so that the effective use of existing tools is urgently needed to combat tuberculosis.

In this study, we focus on cycloserine, an agent that would be added into the initial treatment regimen in priority for MDR-TB according to WHO recommendation, because the evaluation of this drug is greatly hampered by the lack of randomized controlled trials or cohort studies with a reliable outcome measure. To our knowledge, this is the first controlled study to seek to define or optimise the role of cycloserine in drug-resistant tuberculosis treatment. Our data reported an overall treatment success rate of 69.4% within 24 months in the patients treated with cycloserine. Previous studies suggested that the successful outcome rate ranged from 67.5% to 77.0%(18)(9)(19), almost in accordance with our results. There are several possible reasons to explain the slight differences. Firstly, the definition of treatment outcome has been updated and further emphasized the tolerability of the regimens which was likely to be underappreciated before. Secondly, some studies combined adjustive therapy like surgical resection that resulted in improved treatment outcomes(9)(20). Moreover, the accelerated development of pre-XDR-TB and XDR-TB probably reduces the treatment success rate.

Furthermore, compared with the patients in the non-cycloserine group whose regimens mainly included PAS instead of cycloserine, it is suggesting a significant trending towards improved proportion of favorable treatment outcome after the introduction of cycloserine. Based on the current definition we applied, favorable treatment outcome should meet at least three requirements including sputum culture conversion within six months, no sputum reversion, and no severe adverse drug reactions requiring two or more drugs to be discontinued, which indicates that treatment outcome assessments need to integrate efficacy end-points and safety end-points.

Efficacy end-points in this study were mainly measured by time to and proportion of sputum culture conversion. Unlikely drug-susceptible tuberculosis, failure to sputum conversion rather than relapse or sputum reversion accounted for a great proportion of treatment failure(21), suggesting that the current regimens for MDR-TB might not show the strong sterilizing activity. As for cycloserine, our study did not provide sufficient evidence that cycloserine could confer a benefit to culture conversion compared with other standardised treatment regimens. A possible explanation for these results may be that treatment of MDR-TB includes multiple drugs and an observational study without strict placebo controls hardly assesses the efficacy of a single agent. Furthermore, a recent study (22) has showed that more than half of patients with the recommended dosage of 10mg/kg of cycloserine prescription had peak serum concentrations lower than the minimum inhibitory concentrations of the strains isolated from the corresponding patients, suggesting the personal need for adjusting dosages depending on the clinical pharmacokinetic and pharmacodynamic assessments(23).

Adverse drug reaction remains problematic during the treatment course for MDR-TB patients. By contrast with other anti-TB agents, ADR attributed to cycloserine was relatively uncommon with a frequency of 11.1%. Therefore, cycloserine might be regarded as a safer alternative agent to those with frequent severe side effects that tend to result in drug discontinuation and eventually unfavorable clinical outcomes. Similar results were found in a meta-analysis that estimated the frequencies of any ADR from cycloserine at 9.1% (95%CI: 6.4-11.7)(24). Neuropsychiatric reactions, as expect, were representative of adverse effects of cycloserine since its central active mechanism as a partial NMDA-agonist and high brain-blood-barrier permeability(25). In this study, headache was one of the most common side-effects of cycloserine reported by patients although almost headache resolved quickly with an adjustment in the dose or the temporary discontinuation of cycloserine. Seizure was rare, mainly associated with high dosages (especially with concentrations levels exceeding 40 μg/ml)(26), co-administration of fluoroquinolones, and alcoholism(25), but all led to the withdrawal of cycloserine in this study. Psychiatric disturbances were also described on rare occasions in our study but were more complicated to manage for clinicians. In detail, depression or anxiety might be partly attributable to the inadequate social support and lacking of confidence owing to previous poor treatment outcomes, such as the patient (P89) of depression who had been infected with *Mycobacterium Tuberculosis* for more than eight years complicated with post-tuberculosis destroyed lung and complained of unbearable arthralgia during the treatment. The major challenge is the lack of reference standard against which to evaluate psychiatric events. However, the current psychiatric reactions to cycloserine is mainly based on case reports (27) and further controlled studies are needed to validate it scientifically.

To explore the role of cycloserine for patients with different resistance patterns, we did a subgroup analysis and observed the significant improvement in treatment outcomes related to cycloserine in simple MDR-TB patients, which was hindered greatly in complicated MDR-TB patients. The findings from subgroup analyses suggested cycloserine alone are of less benefit without more effectiveness drugs as linezolid for complicated MDR-TB patients(13), indicating the requirement for reprioritization of cycloserine when managing highly resistant forms of tuberculosis. The treatment of complicated MDR-TB, especially XDR-TB, has been considered as a conundrum facing clinicians. In accordance, this study identified the limited beneficial impact of the standardised five-drug regimen in complicated MDR-TB, calling for new or repurposed agents to effectively eradicate extensive drug-resistant *Mycobacterium Tuberculosis*.

This retrospective study has several limitations. First, the major limitations derive from the observational study design which precluded us from controlling of confounding bias well and looking at some important topics, especially pharmacokinetic and pharmacodynamic assessments of cycloserine. Secondly, some strains isolated from patients were missing and thus we did not perform the drug susceptibility testing to cycloserine. Moreover, as cycloserine had not been approved in China until 2014, its availability and affordability require further evaluation.

To summarize, introduction of cycloserine improved the overall favorable outcome of MDR-TB patients. Cycloserine is considered a better-tolerated agent with infrequent adverse side-effects characterized with neuropsychiatric reactions. For simple MDR-TB patients, we believe our results support the use of cycloserine in the setting of correct patient assessment and monitoring. And for complicated MDR-TB patients, more effective treatment options may be considered.

## Acknowledgements

We thank all patients for affording their treatment profiles and all health care workers who participated in this effort.

## Financials

This study is funded by Zhejiang-National Committee of Health and Family Planning Co-Sponsored Project (WKJ-ZJ-07).

